# Amending published articles: time to rethink retractions and corrections?

**DOI:** 10.1101/118356

**Authors:** Virginia Barbour (COPE and QUT), Theo Bloom (BMJ), Jennifer Lin (Crossref), Elizabeth Moylan (BioMed Central) on behalf of COPE working group

**Author notes:** Note: this document does not necessarily represent the views of the organizations listed here.

## Abstract

Academic publishing is evolving and our current system of correcting research post-publication is failing, both ideologically and practically. It does not encourage researchers to engage in consistent post-publication changes. Worse yet, post-publication ‘updates’ are misconstrued as punishments or admissions of guilt. We propose a different model that publishers of research can apply to the content they publish, ensuring that any post-publication amendments are seamless, transparent and propagated to all the countless places online where descriptions of research appear. At the center, the neutral term “amendment” describes all forms of post-publication change to an article. We lay out a straightforward and consistent process that applies to each of the three types of amendments: insubstantial, substantial, and complete. This proposed system supports the dynamic nature of the research process itself as researchers continue to refine or extend the work, removing the emotive climate particularly associated with retractions and corrections to published work. It allows researchers to cite and share the correct versions of articles with certainty, and for decision makers to have access to the most up to date information.

## Introduction

Academic publishing is evolving. It is no longer the case that once published, articles remain unchanged for ever. It is also no longer the case that the final published version is the only version that is made public as depicted in the traditional view of publishing. Increasingly, preprints, datasets and authors' accepted versions (and revised versions), are made available via a variety of mechanisms. A key question that needs to be addressed in the context of this evolving landscape is: are we well-served by the notion of a version of record that is *static* post-publication?

This document sets out the current “best practice” for amending published articles and discusses the problems that are encountered as a result. We suggest a new system which challenges current thinking but proposes a future solution.

The main guideline for handling retractions is the Committee on Publication Ethics (COPE) Retraction Guidelines [1], published in 2009. COPE is a global, multidisciplinary membership organisation of more than 11,000 journals and publishers, which is now in its 20th year. It has evolved into a key body that editors and publishers turn to for advice on many complex publication ethics issues, including on how to handle retractions. Its Retraction Guidelines have formed the basis for the guidelines of many journals, publishers and other publishing organisations. Although these guidelines have been helpful, and were especially relevant in the era of print-based publishing, their consistent implementation has proved more difficult as publishing has evolved. Nonetheless, the guidelines, combined with regular discussion between editors, have provided a core framework for handling retractions that now covers many disciplines and countries. Of particular importance has been the repeated assertion of the overarching intention of the guidelines to assist in correction of the literature, whatever the cause.

Many people are discussing changes in publishing. We bring to this our diverse and collective experience of a number of traditional and start-up publishers as well as of developing infrastructures for open access and other publishing innovations.

## Current best practices

The traditional publishing workflow was originally established to facilitate proper scholarly review within a print publishing paradigm (Figure 1). This process is deeply enshrined in the scholarly communications process, and is carried out with very little variation amongst conventional publishers.

**Figure 1.**
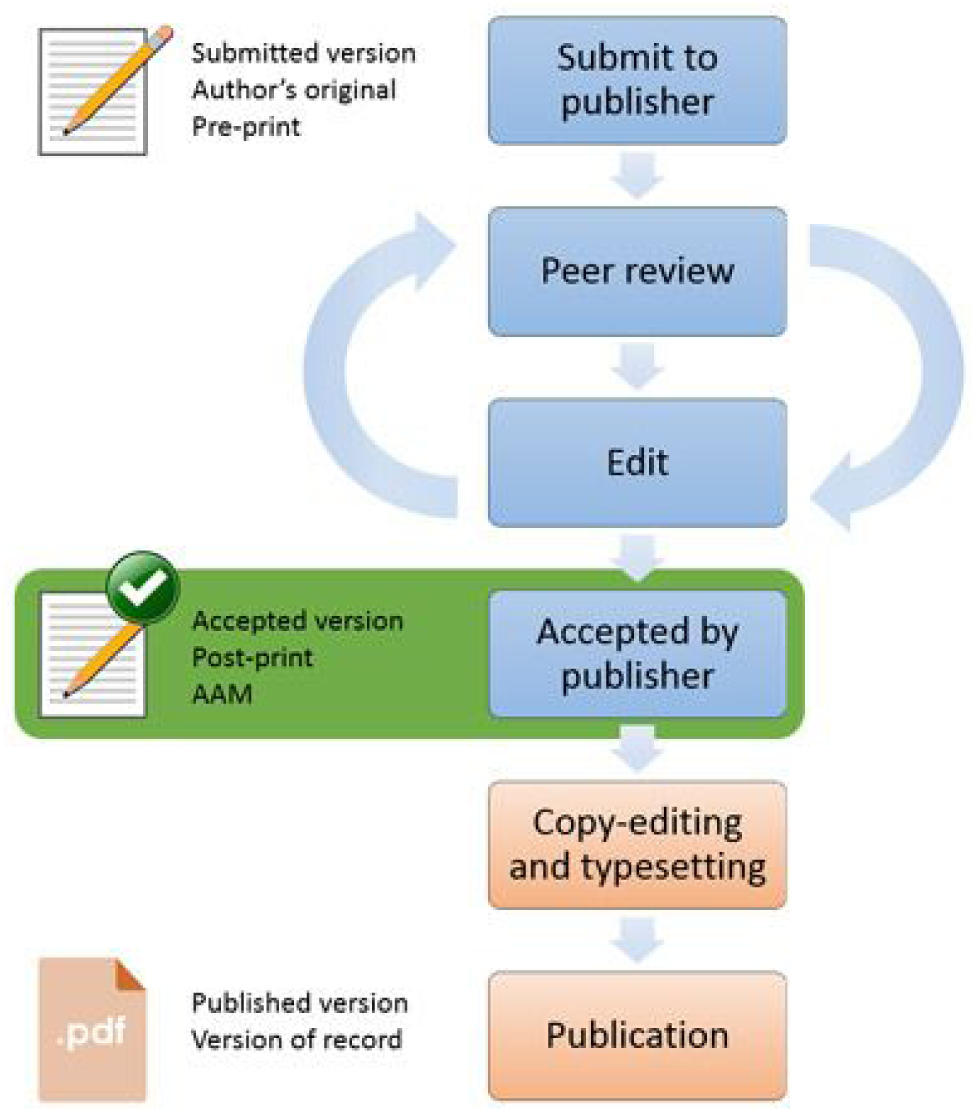
Publishing outline from HEFCE http://www.hefce.ac.uk/rsrch/oa/FAQ/.

Following publication, however, the traditional scholarly communications process gives way to “best practices.” For concerns raised about an article, or author-initiated changes, there are a number of approaches that can be taken to correcting an article, depending on the scale of the issue. Small errors which do not undermine the findings of the published article can be resolved with a “correction”. Historically, some printed journals used to distinguish between errata and corrigenda, according to whether the author or the journal introduced the error ‐ a now meaningless and poorly understood distinction. In other situations it has become more common recently that a comment, editorial or blog (or sometimes many tweets and blogs [2]) may be helpful in providing commentary with or without a correction to the article itself. Letters to the editors also have a long tradition as a place for signed criticism, (e.g. *The BMJ*), though many journals do not allow letters. PubMed Commons [3] now offers a place for any qualified individual to comment on any article that is indexed in PubMed.

Where an article is so seriously flawed or erroneous that the findings can no longer be relied on, then the method of correction is typically wholesale i.e. the article is retracted. COPE guidelines on retraction [1] advise retracting articles if the main findings are found to be unreliable, redundant, plagiarised or if the authors have reported unethical research or failed to disclose a major competing interest which could influence the interpretation of the article. COPE’s intention was to offer practical guidance and not be overly prescriptive (for example the guidelines deliberately did not contain information about the process of retraction and the wording to be used). The guidelines also do not offer guidance on what is to be done after a retraction. For example, some publishers are now experimenting with retracting and replacing an article in its entirety, for example, “retract and replace” by the JAMA network [4]. In other situations, where it is unclear whether a retraction is the final outcome, “Expressions of Concern” typically flag issues that do not yet have a final resolution [1,5,4].

These approaches were designed to help to resolve issues with published articles while maintaining the integrity of the research literature ‐ preserving the original article for the record. But increasingly, these approaches are inconsistently adopted by researchers and editors because many of these mechanisms seem a less than perfect response to an evolving literature in the digital age.

### A fundamental underlying problem

A lack of willingness to engage in proper post publication correction and amendment of the literature is further exacerbated when any type of post-publication “updates” are misconstrued as punishments or admissions of guilt. This is particularly the case with retractions, which many feel has come to be a term loaded with blame and recrimination. It’s fair to say that no one who has had been involved with the retraction of an article – either as an editor, publisher, reviewer or author ‐ has ever walked away from the process feeling wholeheartedly good about the experience. This is even the case if a retraction is done for the best of reasons – a genuine, no fault mistake [6, 7]. As a result, the current system will never be fully embraced as a positive outcome by researchers.

There is a fundamental misconception that retractions are “bad” without pausing to ask *why* the retraction took place. Neutral terminology for a method of correcting published work that implies no fault on any party is therefore needed. In order to provide this, in our view it is important to distinguish the correction of the published record from any investigation or description of misconduct that has occurred. If misconduct or fraud has occurred, this should be reported on, but such reporting should be considered as distinct from the process of correcting the literature. Such a separation is especially important as those who are likely to be responsible for an investigation will be different from those issuing any correction of the published record, and the two may need to happen on quite different time frames. Thus, there may be considerable time spent investigating misconduct and assigning blame. During this time, we feel it is important to alert readers to the possible issues with the published work, and to update the literature without awaiting the final outcome of a lengthy investigation [8].

Although corrections may not be as universally disliked as retractions, once an article has more than one or two corrections, in the current publishing system it is difficult to track what has happened, and this can lead to a feeling of unease from readers and editors. Tracking corrections is even more challenging when they are documented outside of the publication itself, and tracking these external corrections is arguably even more crucial as articles as well as references to them now increasingly propagate across the internet and often do not link back to one version of record (though this has been improved by the development of Crossmark [9]). Furthermore, there are no universal guidelines for corrections, and editors and publishers often act on a case-by-case basis. Many editors and publishers struggle with the need for of a correction notice for a very minor amendment to an article (such as a typographical error that has no effect on meaning), while many readers feel that all amendments post-publication need a clear audit trail.

**Table 1.**
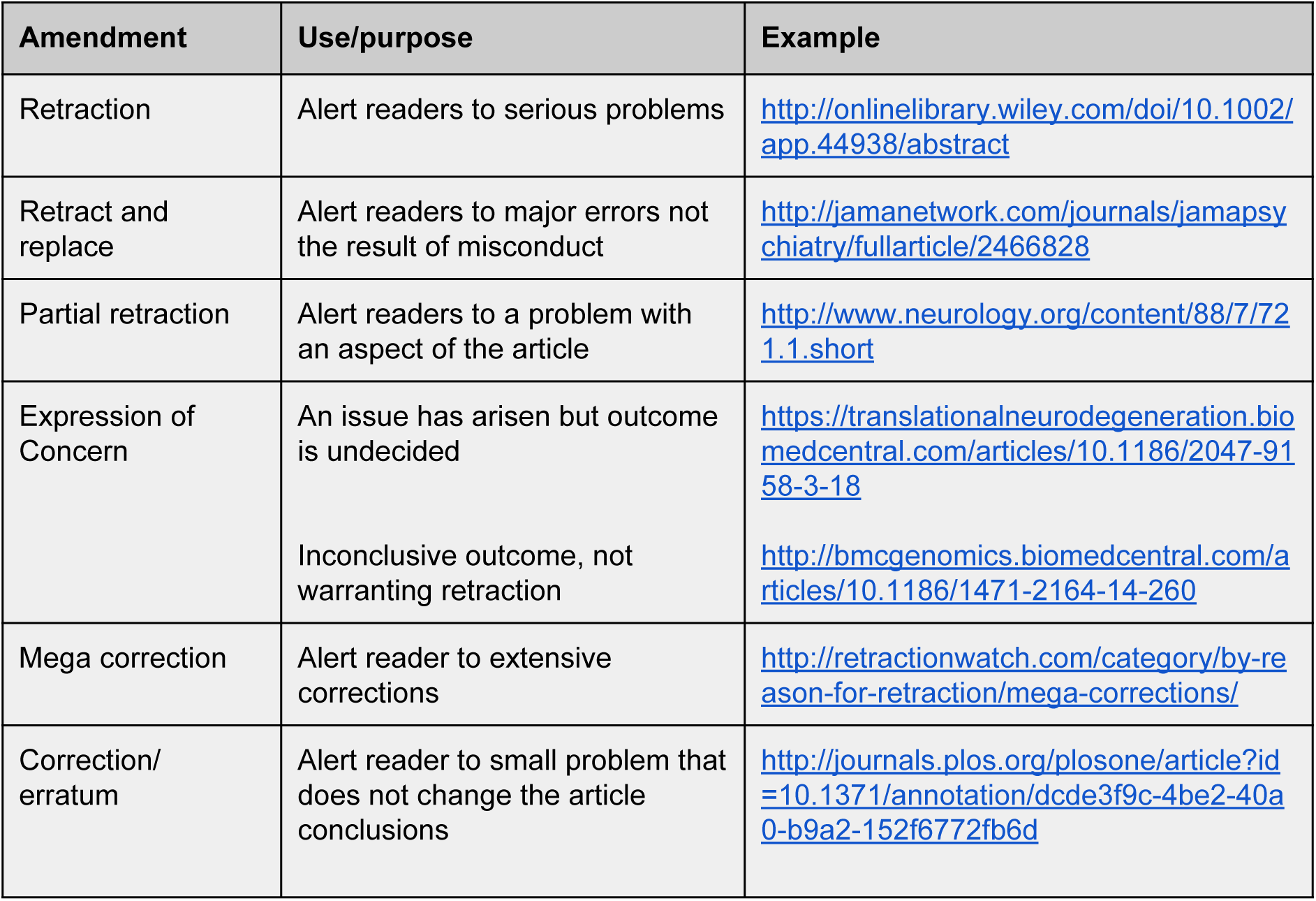
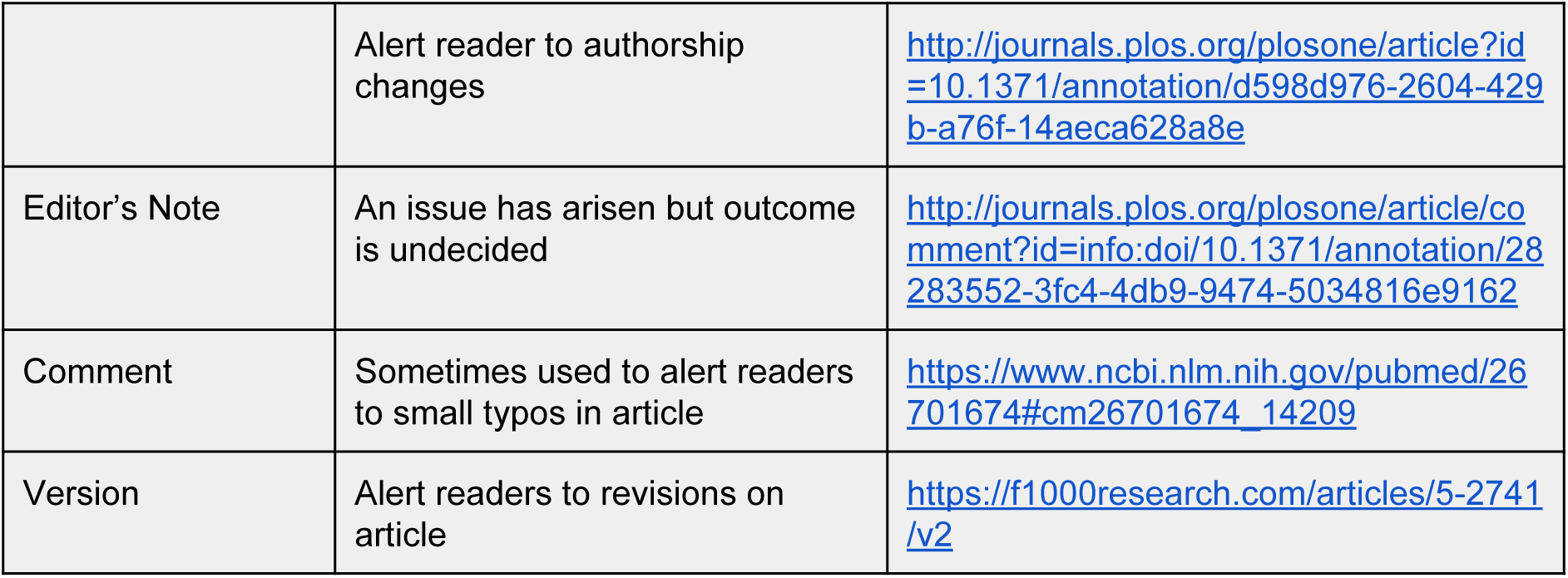
Common amendments in current use.

### Now is the time for change

There are other approaches that can be taken. Outside published research articles, examples from newspapers and blogs take a simple approach to corrections which are, effective, speedy and user-friendly. However, these articles rarely require any cross referencing to external articles and hence the process is much simpler than for academic research articles.

We reaffirm the importance of preserving the integrity of the published literature but we affirm equally strongly that research outputs are now dynamic objects online. The idea of the journal article as a monolithic object that will stand for all time unless formally retracted has gone. Rather we are seeing calls for articles to be viewed as organic publications or “living articles” [9]. If this idea is to be accepted, it is critical to ensure that updates to the scholarly record of a publication are appropriately made and that they properly link the latest update to the original record. Readers need a full trail of changes to an article. We now have the technical tools at hand to do this, and indeed a number of publishers now publish successive versions of works, and it is possible that multiple versions can be preserved for example, if there is a change in crediting or interpretation of the work. These versions can be elegantly handled for readers online by making it clear which version is being displayed and by employing links to help readers navigate between versions (for example, *F1000Research*). Crossmark also provides a vital insurance mechanism, as it displays the publication history consistently across publishers (see Appendix).

We now also have the technical tools to ensure there is clear notation of version history in the citation record. The digital object identifier (DOI) is central to this as a unique alphanumeric string that identifies the content and provides a persistent link to the location of each resource on the Internet. It is an actionable identifier as it resolves to (i.e., takes the user to) the corresponding resource online. It is also descriptive as it binds the DOI to specific metadata about the digital object.

To support versioning, publishers could assign a new DOI or, more logically a DOI with a suffix, to each version, and link to the previous version in the metadata record. Crossref ensures that all versions are linked through the relationship metadata included in the record. Once deposited with Crossref, all versions could be threaded together through their DOIs and made available to systems across the research ecosystem [11]. (Details for how this works are in the next section, below.) This practice provides specificity and precision to the citation record that has not been possible before. Researchers can thus cite a specific version, rather than, unintentionally, the original one. Essentially, we can now, technologically, think beyond the article of record to a number of versions, whether they are preprints or postprints or institutional copies. The primary challenge, however, is publisher (and perhaps institutional) adoption and implementation of practice to deposit and update metadata records as changes occur.

If a fundamental change in how we amend published articles is to be successful, we need both the technology to make it happen (as outlined above) and the will and support from the community to embrace the change.

## A proposal for the future

We propose a future with a fully seamless means of publishing that starts with protocols and registered experiments then moves to results publication and finally onto revisions, with version control. This system incorporates easy corrections, which are themselves integrated with the articles they correct; articles may have multiple versions. “No fault” corrections will be enabled and encouraged– i.e. corrections will be separated from the reason they happened. The degree of reliability of a study will be separated from the notion that the author and/or a prestigious journal provides an absolute guarantee for the work. In all cases by default a reader will see the most recent and up to date version, but they will also be able to navigate to previous versions. There will be full disclosure of publication history and metadata that is made freely available to humans and machines applications.

### The amendment model

In order to provide the widest possible route to a more reliable and useful scientific literature in line with the vision above, we therefore propose the following components of a change to a published article, to be published by the same journal (or other publishing venue) that published by the original research and to be linked to the original article. We propose removing the term retraction altogether and instead using the term “amendment” to describe all forms of post-publication change to an articles.

The term “amendment” carries a neutral tone and is generic enough to apply to a wide array of cases, including the smallest instances such as a typographical error all the way to large retractions. By employing a uniform term, we hope to remove any associated stigma in the context of scholarly literature. When readers encounter each amendment, they read the notice for details on each case, and can judge the article and its revisions as appropriate.

We considered alternative names such as “update” but we felt they imply progress or addition. In particular we strongly feel that retaining the word “retraction” even for the most egregious instances of scientific malpractice would further perpetuate the problem of stigma and is thus not desirable.

We propose an amendment model that includes the following:

1) **Declaration**: Each amendment would make a declaration at the forefront of the document stating that: “The authors and/or the journal wish to make the following amendments to the following published article [article full reference].”
2) **Types of amendment** (based on scale and effect on the key messages)

a) Insubstantial: Examples of this type would be typographical errors, an author-order switch, or other minor amendment to the content or metadata that has no effect on the substance of the article.
b) Substantial: This would cover the types of issues currently worded as Corrections, but might also include clarifications and addenda (which are not currently easy to make under most publishing workflows). Amendments of this type would make changes to one or two small parts of the article but not its whole message. Examples are change of authorship, correction of one figure or method, or addition of a small amount of additional evidence or discussion.
c) Wholesale or complete: the article as a whole is considered unreliable in its current form. There may be elements that remain “correct” but large proportions are not. Instead of “retract and replace” as currently practiced by some publishers, we would recommend “retract and republish” with a new DOI that lands on the newer version and makes plain the chain of events. In the case where authors and/or journal may wish to dissociate themselves from it completely, this is fully noted with a full description in the associated narrative and no attempt to insert new text or other content.
3) **Elements of the amendment**: Every amendment notice would include the following:

a) Who is issuing the notice (an author, all authors, Editor, Publisher, institution), and whether any of this group dissents from the notice. The CReDIT taxonomy may be used to specify the role entailed in crafting the amendment.
b) The scale of amendment, as above
c) Link to the publication it is amending as well as other relevant links to associated resources
d) Date
e) Associated narrative (optional for minor corrections). This is particularly critical in the case of withdrawal of an article without replacement: there needs to be some narrative notice that indicates the reasons.

The process of the amendment within the publication lifecycle is straightforward and consistent for all types of amendment. Whether the incident at hand merits a minor, significant, or wholesale amendment, the publisher can issue the notice, assign a new DOI to it and register it with Crossref, linked to the target publication. Moreover, the same process is also consistent and streamlined to apply at a higher level to include all the various instantiations of the publication from the original publication to the posting of amendments, even the publication of subsequent versions of the paper (Figure 2).

**Figure 2.**
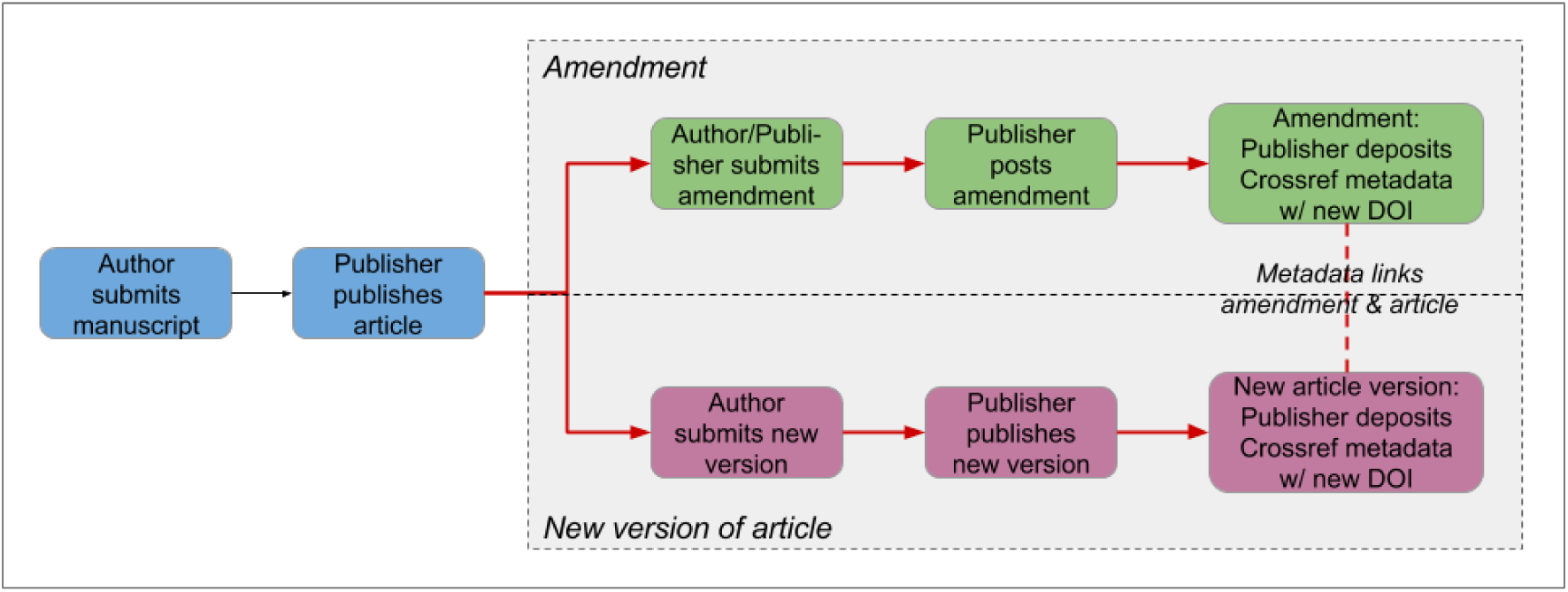
The amendment process.

This version of scholarly communications supports the dynamic nature of the research process itself as researchers continue to refine or extend the work. They can publish updates along the way, sharing out the latest findings, analysis, and conclusion. For each version of a paper, an amendment can be issued. In each case, the publisher assigns a new DOI to each of the pieces and deposits the metadata with Crossref so that researchers can cite with clarity and specificity.

### Applying the amendment model: an illustration

Models are generalized abstractions, and in this case the proverbial saying “the devil is in the details” thoroughly applies. To illustrate the proposed amendment model, we apply the model to a case that contains elements of a number of COPE cases. The underlying issue relates to a duplicated gel image.

An institution found misconduct from one of their senior scientists who reused the same images to show controls in many figures. Once the publisher was notified, they opened an investigation on the three publications found to be affected. After consulting with the original referees, the publisher gave the authors an opportunity to publish corrections with corrected images. The authors provided new control images and corresponding results for most, but not all, of the figures in question. The original referee determined that the corrections sufficed, and that it was not necessary to repeat all the experiments. An editorial board member, however, still felt that the three articles should be retracted due to fraudulent behaviour.

The publisher identified three options for proceeding with this incident:

1. Retract all three articles because the authors have lost credibility and misconduct was confirmed in the duplication of control images.
2. Publish a correction notice on each of the articles to warn the readers about the duplication of control images. While issued as a “correction”, the text would amount to an “expression of concern”.
3. Publish the corrections supplied by the authors.

COPE’s recommendation would be for the publisher to apply the findings of the institution’s investigation to decide between options (1) or (2), but the institution would not share it. The publisher decided to retract the two articles in which the questioned experiments were not repeated, and publish a correction for the article where the authors provided new results and controls.

The publisher was conscientious and thorough, demonstrating responsible stewardship with its content˙. In the years that this case, as is typical with many of these cases took to resolve, many of the publisher’s discussions with reviewers and editorial board centred around the application of a retraction or correction designation. In the amendment model proposed, these exchanges are no longer necessary for an amendment to be implemented. Once the gel image duplication is identified and due diligence applied, the publisher can expediently document the findings and communicate this to the research community, specifically to the readers of the three publications. The question of the author’s motive, personal credibility, and trustworthiness, while relevant in other contexts, did not need consideration in this incident where the publisher was acting as editorial steward for the research results they disseminate. The model proposed thus has the potential for resolving the post-publication issues more expediently, by focusing the resolution process on the publisher role and aim at hand: to communicate the latest status of the research findings included in the publication. Readers would be much better served by a rapid explanation of the issues, rather than there being no notification during the necessary “due process” of a misconduct investigation.

If the publisher were to have applied the amendment model to its process, the publisher would have been able to focus on the issues relevant to issuing a notice and communicate to their community of researchers much faster. This approach does not compromise the thoroughness of any deliberations regarding status of the publication itself, but rather winnows out the more complicated discussions that lie outside of the publisher’s immediate realm, such as research misconduct in general, institutional compliance, misconduct regulations, and any other actions related to misconduct that might have occurred.

**Figure 3.**
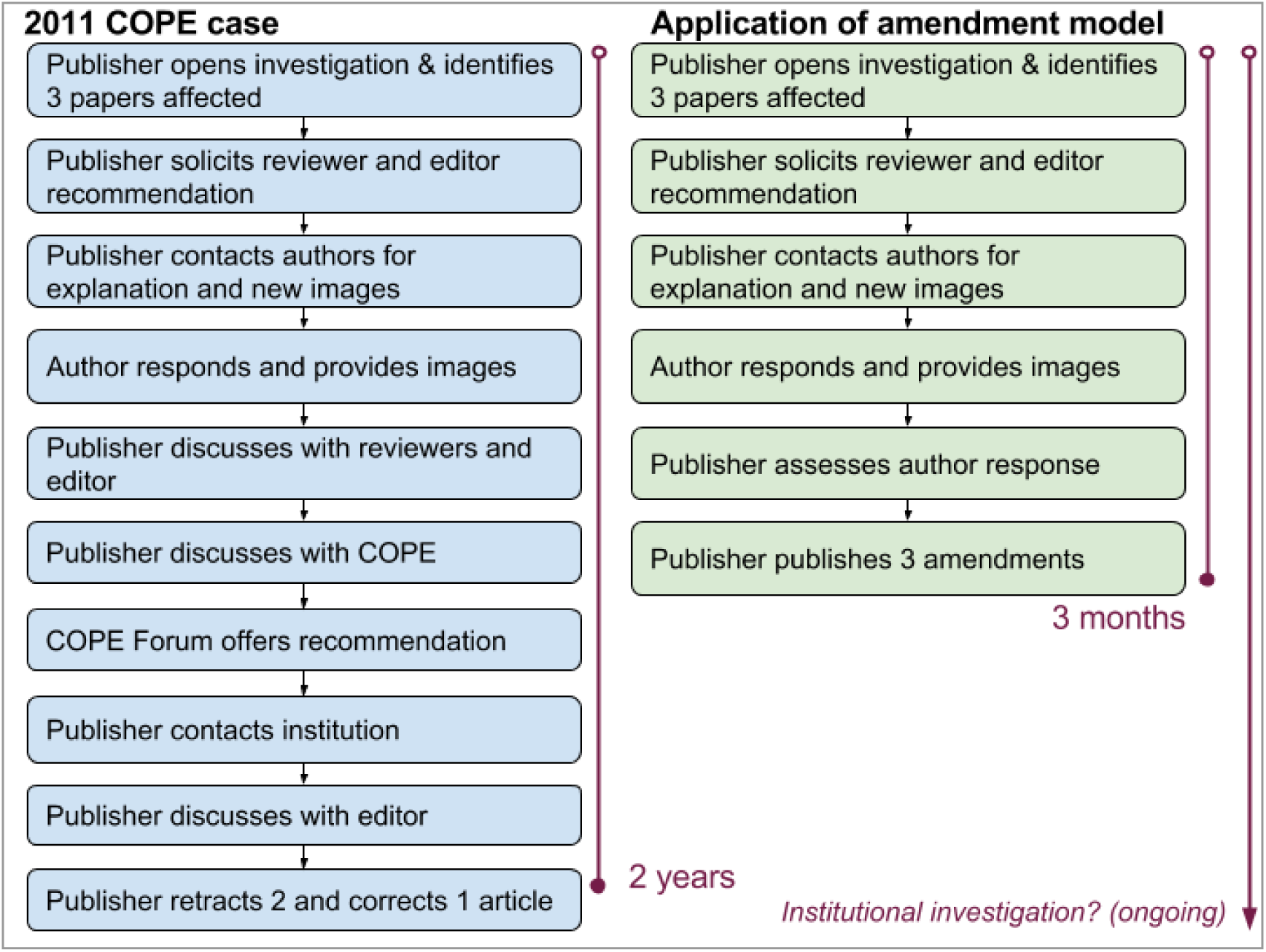
COPE example & application of amendment model.

### Amendment display & linking

While this amendment model simplifies the process by which published results are shared and updated, it also increases the potential number of components that might be published from a single set of research results. As such, linking amendments to their associated articles and across individual versions needs to be carefully implemented online so readers can easily navigate to the research results intended.

Since every publisher employs their own specific design approaches to content delivery, we recommend the following linking and display strategies to ensure that amendment display fully supports editorial intent:

1. **Notice and article**: Link from each amendment to the respective article it amends and vice versa, so that the reader easily navigate back and forth between the notice and the research itself.
2. **Article versioning**: Each article version has its own DOI and URL, which persists even with the publication of subsequent versions. Where the reader is on a dated version, clearly communicate this date on the article. In every single version, provide a clean and simple way for readers to navigate between versions. In cases where the publisher is linking to the article in general (e.g. from a journal home page, etc.), directing users to the latest version is recommended.
3. **General amendments to article version:** The amendment notice links back to the specific article version to which it amends. Amendment #1 can link to Article Version 1. Amendment #2 might also link to Article Version 1 if a subsequent one is issued about the same article version. If Amendment #3 is issued for the second version, then it links to Article Version 2.
4. **Figure legend amendment:** In the event of a correction to a figure legend, the link should direct users to the latest version of the article, complete with the correct figure legend. Readers can then go back and look at it with the wrong legend as they wish (i.e. be transparent about the change). This is akin to the journalistic model and is much cleaner.
5. **DOI construction:** Although any two DOIs can be linked via CrossRef to explain the relationship between versions of an article, or between an article and an amendment, we suggest that the relationship between these would be more obvious to human readers if the original article’s DOI were given a suffix (e.g. doi.org/10.1136/bmj.j1072.1; doi.org/10.1136/bmj.j1072.2 and so on)
6. **DOI resolution**: The DOI should direct users directly to the specific document at hand, whether that is Amendment 1, Article Version 1, Amendment 2, Article Version 2, etc.

### Amendment metadata & propagation

Minor amendments must be evident to readers but may be reasonably handled in the way blogs and news outlets handle such minor changes provided that there is a technological option to allow for recording that there has been a minor change since publication,which is propagated to systems that index articles. Significant amendments must be evident to readers but also to machine harvesters of the literature, and must be inextricably and permanently linked to the original article. Some instances of the third type of amendment would mean publication of a new article, with a new DOI linked to the original article.

For publication updates to reach all the places online where articles are read, indexed, shared, discussed, recommended, etc. publishers need to make the metadata available in a central store where the data is freely available for systems and applications to consume. Crossref currently provides this facility and their metadata framework fully supports full disclosure of amendments. Once metadata updates are available from publishers, systems need to apply the latest information wherever the publications affected may appear online. All metadata are openly licensed for reuse, propagated through a variety of interfaces and formats (Crossref APIs).

Anyone can search the scholarly registry (humans and machines) to get the latest updates for any publication regardless of origin in the Crossref corpus (85+ million publications at time of writing). Publishers can flag this not only in their own content (online and PDF versions) via Crossmark, but also in the references of papers they publish. They can propagate these notices through other delivery channels offered such as eTocs, RSS feeds, recommendations, etc. Non-publisher platforms such as indexers, reference managers, recommendation systems, social bookmarking tools, researcher profile systems, etc. can apply the update information and bibliographic metadata to the content they display as well. This information is also potentially useful for research information systems used by funders and research institutions, which also track scholarly outputs.

## Conclusion

Our current system of correcting research post publication is failing both ideologically and practically. We propose a model that publishers of research can apply to the content they publish which ensures that any post-publication amendments are seamless, transparent and exposed and propagated to all the countless places online where descriptions of research appear. We believe that this proposal, will allow us to have a system which both incorporates new technological thinking and removes the emotive climate now associated with retractions and corrections to published work. It also exploits the opportunities of new technologies to allow researchers to cite and share the correct versions of articles with certainty and for decision makers to have the most up to date information in order to support the research enterprise.

## Declarations

### Funding

There was no funding for this project which has been conducted on a voluntary basis.

### Competing Interests

Virginia Barbour and Theodora Bloom are both on the Eighth International Congress on Peer Review and Scientific Publication Advisory Board. Virginia Barbour is the Chair of COPE. Elizabeth Moylan is on the COPE council.

### Author contributions

All authors contributed equally and are listed in alphabetical order. This work was conducted as part of an ongoing open discussion within an initial working group and a wider consultation.

## Acknowledgments

We are grateful for comments and feedback from Geoffrey Bilder, Peter Doshi, Andy Collings, Michaela Torkar and Liz Wager.

## Appendix A ‐ Crossmark

The Crossmark identification service from Crossref sends a signal to researchers that publishers are committed to maintaining scholarly content [9]. It gives scholars the information they need to verify that they are using the most recent and reliable versions of a document. Readers simply click on the CrossMark logos on PDF or HTML documents, and a status box tells them if the document is current or if updates are available. Clicking on a Crossmark logo may also provide important publication record information about the document. This information, provided at the option of the publisher, might include peer review, publication history, funding sources, location and links to data sources, Similarity Check plagiarism screening, or rights. It notifies readers of changes to content in a consistent way, regardless of who publishes it or where on the web it is stored. Crossmark also encourages users to go to the publisher’s site for the verified, up-to-date copy.

Crossmark also has dedicated support for clinical trials metadata. Publishers add the clinical trial numbers to the Crossref DOI metadata via three new fields: clinical trial number, clinical trial registry where trial is registered, and trial stage (pre-results, results or post-results of the trial). Crossref displays the clinical trial metadata on the respective papers for all participating Crossmark publishers as well as links to all the publications that reference the same clinical trial. Publishers can collect this information upstream and disseminate it using the existing Crossref infrastructure.

